# Splitting and filling the gaps: a reorganization of Corymbiglomeraceae and new taxa from trans-Pacific tropical regions

**DOI:** 10.64898/2026.01.28.702294

**Authors:** Thomas Crossay, Martin Hassan Polo-Marcial, Keyvan Esmaeilzadeh-Salestani, Mariana Bessa de Queiroz, Juliana Luiza Rocha de Lima, Luis Alberto Lara-Pérez, Javier Isaac de la Fuente, Sylwia Szczecińska, Maurice Wong, Leho Tedersoo, Bruno Tomio Goto, Franco Magurno

## Abstract

Diversisporales comprises species with worldwide distribution that produce glomoid, otosporoid, or tricisporoid spores. The recent reorganization of the order by Oehl et al. (2016) recognizes two families, Diversisporaceae and Corymbiglomeraceae, comprising one and five genera, respectively. Several Glomeromycotan specimens collected in northern and southeastern Mexico and in French Polynesian atolls were characterized using both morphological and molecular analyses. Phylogenetic inference revealed that they represent new members in the Diversisporales, supporting the reorganization of the genus *Redeckera* into three independent lineages: *Albocarpum* gen. nov., with *A. arenaceum* sp. nov., *A. leptohyphum* sp. nov., and *A. fulvum* comb. nov., *Pulvinocarpum pulvinatum* gen. et comb. nov., and *Redeckera*, which retains five species, including *R. varelae* sp. nov. In addition, we described *Melanocarpum mexicanum* gen. et sp. nov. and *Diversispora papillosa* sp. nov. A broader phylogeny, based on eDNA sequences and representative of Diversisporales species, including the newly described taxa, further supported the split of *Redeckera* and suggested three additional clades likely corresponding to a new family and two new genera, awaiting the discovery of representative morphospecies to be formally described. Using eDNA sequences metadata, the occurrences of the newly described taxa were mapped, allowing to recognize distribution patterns, mostly in the pantropical zone, distinguish widespread and rare species, and suggest possible endemisms. Finally, the coexistence of species forming large sporocarps (*A. fulvum* and *A. leptohyphum*) alongside species forming spores in loose aggregates (*A. arenaceum*), prompted us to propose a possible sporulation dimorphism in *Albocarpum*, an argument previously raised to explain the nested placement of *Corymbiglomus* and *Paracorymbiglomus* within the *Redeckera* clade.

## Introduction

Diversisporaceae was introduced by Schwarzott et al. (2001) as a putative new family representing the species in the clade GlGrC (*Glomus versiforme*, *G*. *spurcum*, and *G*. sp. W2423), previously included in the Glomaceae (= Glomeraceae). Later, the family was formally validated by Walker and Schüßler (2004), who established *Diversispora*, with *D. spurca* (≡ *Glomus spurcum*) as the type species.

Since its establishment, Diversisporaceae has undergone numerous revisions, including the introduction of new genera, some of which were later considered invalid, and the transfer and reclassification of several species (Oehl et al. 2026). Palenzuela et al. (2008) introduced the genus *Otospora*, with the novel species *O. bareae*, within the Diversisporaceae, based on morphological traits of spore development resembling those of *Acaulospora*, and SSU rDNA sequences. Schüßler and Walker (2010) further enlarged the family with the genus *Redeckera*, typified by *R. megalocarpa* (as *R. megalocarpum* ≡ *G. megalocarpum*), characterized by production of large glomerocarps covered by a peridium, and supported by phylogenetic analyses of the SSU–ITS–LSU nrDNA marker. *Redeckera fulva* (as *R. fulvum* ≡ *G. fulvum*) and *R. pulvinata* (as *R. pulvinatum* ≡ *G. pulvinatum*) were also included. Later, Oehl et al. (2011a), considering the morphological similarities among glomerocarps also transferred *G. avelingiae*, *G. canadensis*, and *G. fragilis* to *Redeckera*. Oehl et al. (2011b) introduced the genus *Tricispora*, reclassifying *Entrophospora nevadensis* as *T. nevadensis*. However, this genus as well as *Otospora*, were later considered synonymous with *Diversispora* by Tedersoo et al. (2024a) due to their position nested in *Diversispora*. Błaszkowski and Chwat (2013) described *Corymbiglomus* based on SSU–ITS–LSU nrDNA analyses, including *C*. *corymbiforme*, *C. globiferum*, and *C. tortuosum*, previously assigned to *Glomus*. Phylogenetic evidence revealed that *C. tortuosum* forms a distinct genus-level clade within Diversisporaceae, prompting the establishment of *Sieverdingia* for *S. tortuosa* (Błaszkowski et al. 2019). Symanczik et al. (2018), relying on SSU–ITS–LSU nrDNA and RPB1 genes phylogenies, showed that *Diversispora omaniana* does not belong to *Diversispora* but instead represents a novel genus, *Desertispora*. Recently, inconsistencies regarding the monophyly of *Redeckera* have been reported in Tedersoo et al. (2024a) and Błaszkowski et al. (2025), which led to the establishment of the new genus *Paracorymbiglomus* to accommodate *C. globiferum* and *C. pacificum*, restricting *Corymbiglomus* to its type species, *C. corymbiforme*.

In a recent review, combining spore-based morphology and phylogenetic analysis of the 45s rDNA sequences, three new orders, five new families and several combinations were introduced in Glomeromycetes. Among them, Diversisporales was split into four orders: Diversisporales, Sacculosporales, Acaulosporales, and Pacisporales. The new family Corymbiglomeraceae was established, including the genera *Desertispora*, *Corymbiglomus*, *Paracorymbiglomus*, *Redeckera*, and *Sieverdingia,* leaving the sister family Diversisporaceae with only *Diversispora* (Oehl et al. 2026).

Different types of spore formation are observed in Diversisporales, including otosporoid (laterally on the persistent neck of a terminal or intercalary sporiferous saccule at some distance from the saccule terminus), tricisporoid (within the hyphal neck of a tightly attached terminal or intercalary sporiferous saccule, closely attached to the saccule terminus) or, most commonly, glomoid-like types (produced terminally on the subtending hypha) (Oehl et al. 2011a, b; Wijayawardene et al. 2025). When glomoid spores present a spore wall that is not continuous with the subtending hyphal wall, they are also referred to as diversisporoid, a morphotype typical of *Diversispora* and *Redeckera* (Oehl et al. 2011b).

The formation of glomoid-like spores in sporocarps represents another feature shared by *Diversispora* and *Redeckera*. However, all known *Redeckera* species consistently produce spores only in compact epigeous or sub-hypogeous glomerocarps with a peridium, whereas *Diversispora* species produce spores singly, in small or large loose clusters, or within epigeous or sub-hypogeous glomerocarps, generally without a peridium (except *D. sporocarpia*) (Jobim et al. 2019a). Other genera in Diversisporales, such as *Desertispora*, *Corymbiglomus*, *Paracorymbiglomus*, and *Sieverdingia* produce hypogeous spores, either singly or in small loose clusters (Symanczik et al. 2018; Błaszkowski et al. 2019, 2022, 2025).

The detection of sporocarps through active searches has led to the discovery of new species, mainly in vegetation types with some degree of conservation (Redecker et al. 2007; Jobim et al. 2019a; Błaszkowski et al. 2021; Yamato et al. 2024). However, there are also recurrent records in disturbed forests by wildfires in Australia (McGee 1986; McGee and Trappe 2002), and in corn monocultures in the Yucatan Peninsula (Guzmán 2003; Polo-Marcial et al. 2021). Most known AMF species have been described based on spores/clusters recovered directly from field soil or cultivated in trap or single-species cultures (Goto et al. 2024). Despite ongoing taxonomic efforts, the rate of descriptions of new Glomeromycota species remains low (Goto et al. 2024). This rate is even lower when describing glomerocarpic species (Jobim et al. 2019a), reflecting the difficulties associated with their detection, cultivation, and biological characterization.

During the analysis of several single-spore isolates and glomerocarps from Mexico and French Polynesia, we identified specimens representing previously undescribed taxa within Diversisporales. Additionally, our results also strengthened earlier suggestions that *Redeckera* is polyphyletic. Accordingly, the aims of this study are to (i) provide detailed morphological descriptions of the newly discovered taxa, (ii) elucidate their phylogenetic placement within the Diversisporales, and (iii) revisit the genus *Redeckera*, utilizing a broad eDNA phylogeny approach to clarify the status of its species.

### Materials and Methods Sampling

Two spore morphotypes (provisionally named species “Fc1” and “Fc2”) were isolated from two atolls of French Polynesia: Rangiroa and Fakarava, both located in the north-western Tuamotu archipelago in the central South Pacific. “Fc1” was collected in Rangiroa (14°58’24.3“S, 147°37’28.4”W), and “Fc2” in Fakarava (16°08’23.8“S, 145°35’26.9”W). The spores were sampled, collected and grown in single-species cultures according to the method proposed by Crossay et al. (2018). The study sites present a mosaic of shrubby autochthone vegetation and agronomic species. There are several unifying features: shrubby vegetation is evergreen, 1–7 m tall, more or less bushy, and most species are sclerophyllous. Most plant species have a well-developed superficial root system that allows them to grow in shallow soil composed mainly of coral debris and sand. The air temperature ranges from 22°C to 31°C. Rainfall is around 1,500 mm/year, with values ranging from 900 to 2,200 mm (Duvat et al. 2020, 2022).

Glomerocarps were isolated from several sites in southeastern and northern Mexico, characterized by different vegetation types. The glomerocarps were collected according to the method proposed by Jobim et al. (2019a). Within the collection, three specimens were selected on the basis of their morphological characteristics and provisionally named specimens (spec.) “R2”, “De” and “Dt”.

Spec. “R2” was isolated from a highly disturbed fragment of the semi-evergreen tropical forest within the Zazil Urban Ecological Park, in the municipality of Othón P. Blanco, Quintana Roo (18°30’25.9“N, 88°19’11.4”W, 6 masl). The forest is characterized by a tree stratum that reaches heights of up to 30 m and a partial foliage loss (25-50%) during the dry season. The annual average temperature and precipitation are 20°C and 1200 mm, respectively. The vegetation is dominated by Fabaceae, Sapotaceae, and Meliaceae (Martínez and Galindo-Leal 2002; Islebe et al. 2015).

Spec. “De” was isolated from a fragment of low deciduous forest from Hermenegildo Galeana, Campeche (18°10’42.7“N, 89°14’30.2”W, 190 masl). The vegetation is dominated by Arecaceae, Burseraceae, Polygonaceae, and Sapotaceae, with annual precipitation of 600–800 mm and an average annual temperature of 25°C (Islebe et al. 2015).

Spec. “Dt” was isolated from the tropical mountain cloud forest of the El Cielo Biosphere Reserve in Gómez Farías, Tamaulipas (23°01’26.2“N, 99°09’57.5”W, 540 masl). The dominant vegetation consists of species belonging to Altingiaceae, Fagaceae, and Rosaceae (Arriaga 2000). Furthermore, glomerocarps of *Redeckera fulva* (“R1”) were isolated directly from the primary root of *Lonchocarpus yucatanensis* Pittier in a preserved fragment of the tropical semi-evergreen forest in Petcacab, municipality of Felipe Carrillo Puerto in Quintana Roo (19°12’02.5“N, 88°26’54.0”W, 30 masl).

In order to obtain monospecific cultures, internal fragments of sporocarps were pulverized, following the method described by Błaszkowski (2012). The spores were kept in distilled water and refrigerated for 24 hours before being inoculated onto *Sorghum* sp. and *Zea mays* L.

### Molecular analysis

Total genomic DNA of the two isolates from French Polynesia was extracted from ca. 30 single spores, by crushing the spores with a micropestle in a vial containing 30 μL of distilled water, followed by centrifugation to pellet the debris. Extraction of genomic DNA from specimens collected in Mexico was performed using the DNeasy PowerSoil Pro kit (Qiagen, Hilden, Germany), as described in Magurno et al. (2024).

Amplicons of SSU–ITS–LSU nrDNA partial genes (thereafter referred to as 45S) were obtained by PCR with the primer pairs SSUmAf–LSUmAr, followed by a nested reaction with SSUmCf–LSUmBr (Krüger et al. 2009), using the Phusion Plus DNA Polymerase (Thermo Fisher Scientific, Waltham, MA, USA) with a universal annealing temperature of 60 °C, according to the producer’s instructions.

PCR products were purified with GeneJET PCR Purification Kit (Thermo Fisher Scientific, Waltham, MA, USA), and then cloned with CloneJET PCR Cloning Kit (Thermo Fisher Scientific, Waltham, MA, USA). After screening under the same conditions as for the nested PCR, plasmids were extracted using the GeneJet Plasmid Miniprep Kit (Thermo Fisher Scientific, Waltham, MA, USA) and sequenced at Genomed S.A. (Warsaw, Poland). Sequences were deposited in GenBank (PX612380-PX612384; PX661504-PX661521; PX700901-PX700907)

### Phylogenetic analyses

Phylogenetic inference of the putative new species was conducted in two stages. First, Maximum likelihood and Bayesian analyses were performed using a dataset of 136 sequences from 36 representative members of Diversisporales that possess the 45S barcode (or part of it), along with the sequences obtained in this study. Members of *Sacculospora* were also included as the outgroup.

The analyses aimed to determine the phylogenetic placement of the putative new species and evaluate their status as distinct lineages. The dataset was aligned with MAFFT v.7 (Katoh and Standley 2013) and E-INS-i as iterative refinement methods (http://mafft.cbrc.jp/alignment/server/). Phylogenetic analyses based on Bayesian inference and maximum likelihood were carried out at the CIPRES Science Gateway 3.1 (Miller et al. 2010), using MrBayes v3.2.7 (Ronquist et al. 2012) and RAxML-NG (Kozlov et al. 2019). Model settings and partitions were configured as in Magurno et al. (2024). For Bayesian analysis, the number of generations was increased to 5 million, with a stop rule at a split frequency standard deviation of 0.01. Phylogenetic trees from the two analyses were visualized, merged, and edited in TreeGraph 2 (Stöver and Müller 2010). Clades were considered supported when Bayesian posterior probabilities were ≥0.95 and ML bootstrap values were ≥70%.

The second stage of analysis involved a broader phylogeny based on 602 eDNA sequences and 83 sequences from morphospecies. The analyses aimed to assess the polyphyletic status of the genus *Redeckera* and detect autonomous supported lineages representing potential new genera.

Three nucleotide sequence repositories – EUKARYOME v.1.9.4 (Tedersoo et al. 2024a), NCBI (Sayers et al. 2024), and UNITE v.9.1 (Abarenkov et al. 2024) were used to download sequence data assigned to Glomeromycota. Sequences labeled as unidentified fungi from NCBI and UNITE were preliminarily assigned to rough taxonomic groups using BLASTn queries against curated references in EUKARYOME v.1.9.4. Those identified as Diversisporales were selected to build a 45S dataset.

Sequence alignment was performed using MAFFT v.7, followed by manual trimming to correct misalignments and remove poorly aligned terminal regions using AliView v1.26 (Larsson 2014). To ensure taxonomic reference, at least one representative sequence from each described species was included to delineate clades and support taxonomic assignments. Subsequent filtering of alignments was performed using ClipKIT v.1.4.0 (Steenwyk et al. 2020) to remove positions lacking phylogenetic information. The initial three rounds of phylogenetic analysis focused on detecting and excluding low-quality and chimeric sequences. In the fourth round, only high-quality reads were retained and subsequently used to construct the final phylogenetic tree.

The tree was reconstructed using maximum-likelihood methods in IQ-TREE v.2.2.5 (Minh et al. 2020). The analysis employed a partitioned dataset (as SSU, ITS1, 5.8S, ITS2, LSU) with a GTR+I+G nucleotide substitution model, including 1000 ultrafast bootstrap replicates and 1000 SH-aLRT tests for branch confidence. The resulting tree was visualised and taxonomically reannotated using FigTree v1.4.5 (Rambaut 2024). Furthermore, metadata available in the EUKARYOME v.1.9.4 database were used to map the occurrences of members of the detected lineages and their distribution across distinct biomes (Tedersoo et al. 2024a).

### Glomerocarp analysis

Macroscopic characteristics such as color, texture, size, shape, presence of rhizomorphs, and aroma were recorded from fresh glomerocarps. In addition, fragments of peridium, gleba and base of glomerocarps were mounted in water, PVLG (lactic acid, polyvinyl alcohol/lactic acid/glycerol), and PVLG + Melzer’s reagent (1:1) v/v. Fragments of each glomerocarp were stored in silica gel to preserve genetic material.

### Microscopy and Nomenclature

Morphological, phenotypic and histochemical characteristics of the spores were characterized from 50–100 spores mounted in water, PVLG, and a mixture of PVLG and Melzer’s reagent (1:1, v/v) (Omar et al. 1979; Błaszkowski 2012). Spore preparation was performed, and micrographs were obtained, following the protocol described by Błaszkowski (2012). Spore wall layer types were defined according to Błaszkowski (2012) and Walker (1983). Color names were used according to Kornerup and Wanscher (1983). We adopted the terms “glomerospores” and “glomerocarps” proposed by Goto and Maia (2006) and Jobim et al. (2019a), respectively. Fungal nomenclature and the authors of fungal names are from the MycoBank (https://www.mycobank.org/). Reference specimens of the proposed new species were deposited as holotypes in the collection of fungi of the XAL Herbarium (Instituto de Ecología, A.C., Mexico) and in the Mycological Herbarium of the National Museum (Paris, France).

## Results

### Molecular data and phylogenetic analysis

Five to six partial 45S sequences were successfully obtained from each of the five specimens analyzed. According to BLASTn comparisons, all specimens differed from the closest described species with a minimum percentage of dissimilarity ranging from 4% to 10.6%.

Two additional sequences from the *Redeckera fulva* specimen “R1” were also obtained since only short sequences for the species were available in GenBank.

Maximum likelihood (ML) and Bayesian (BI) phylogenies (Figure 1) were consistent in the analysis based on the dataset of representative species for Diversisporales.

**Figure 1.**
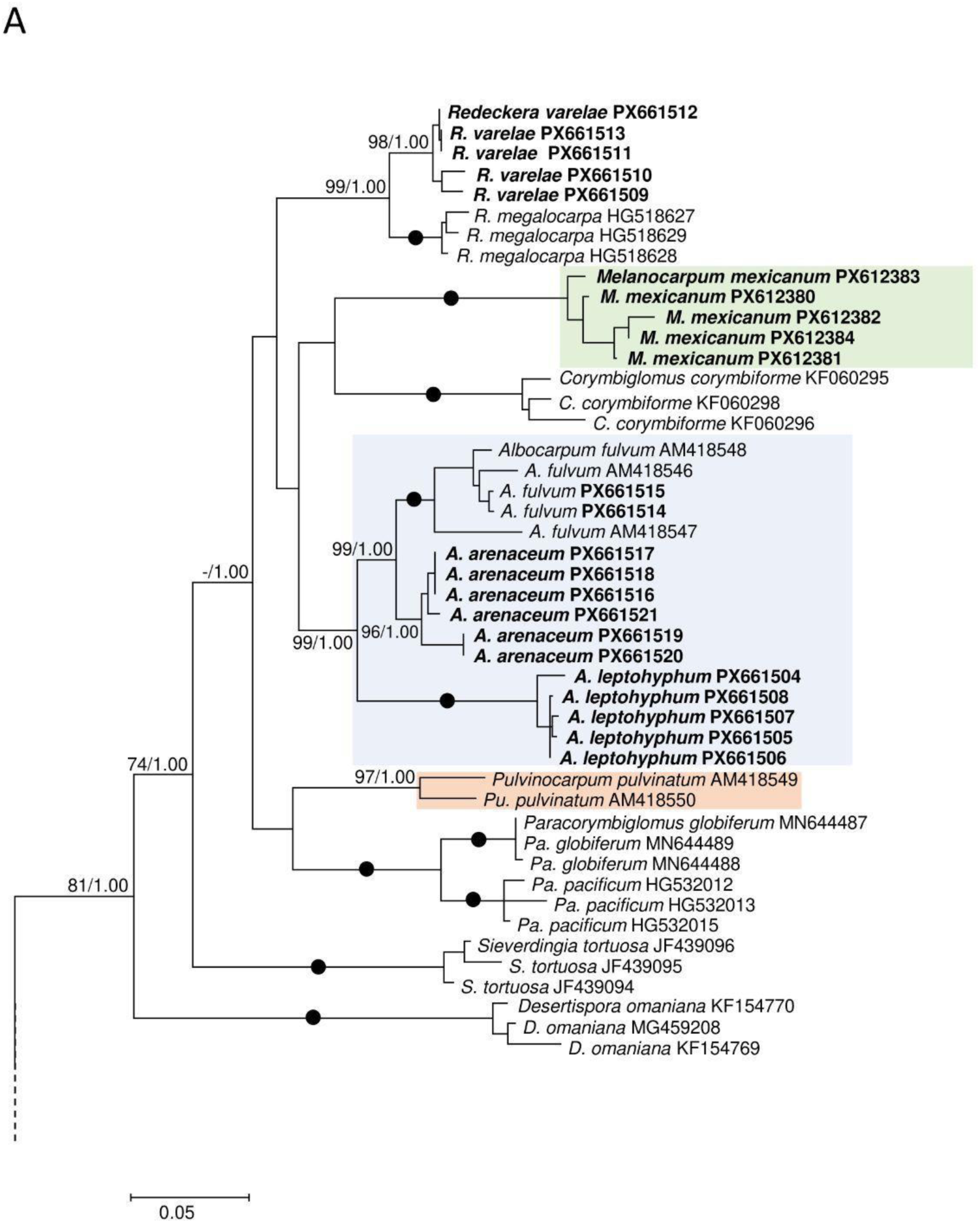
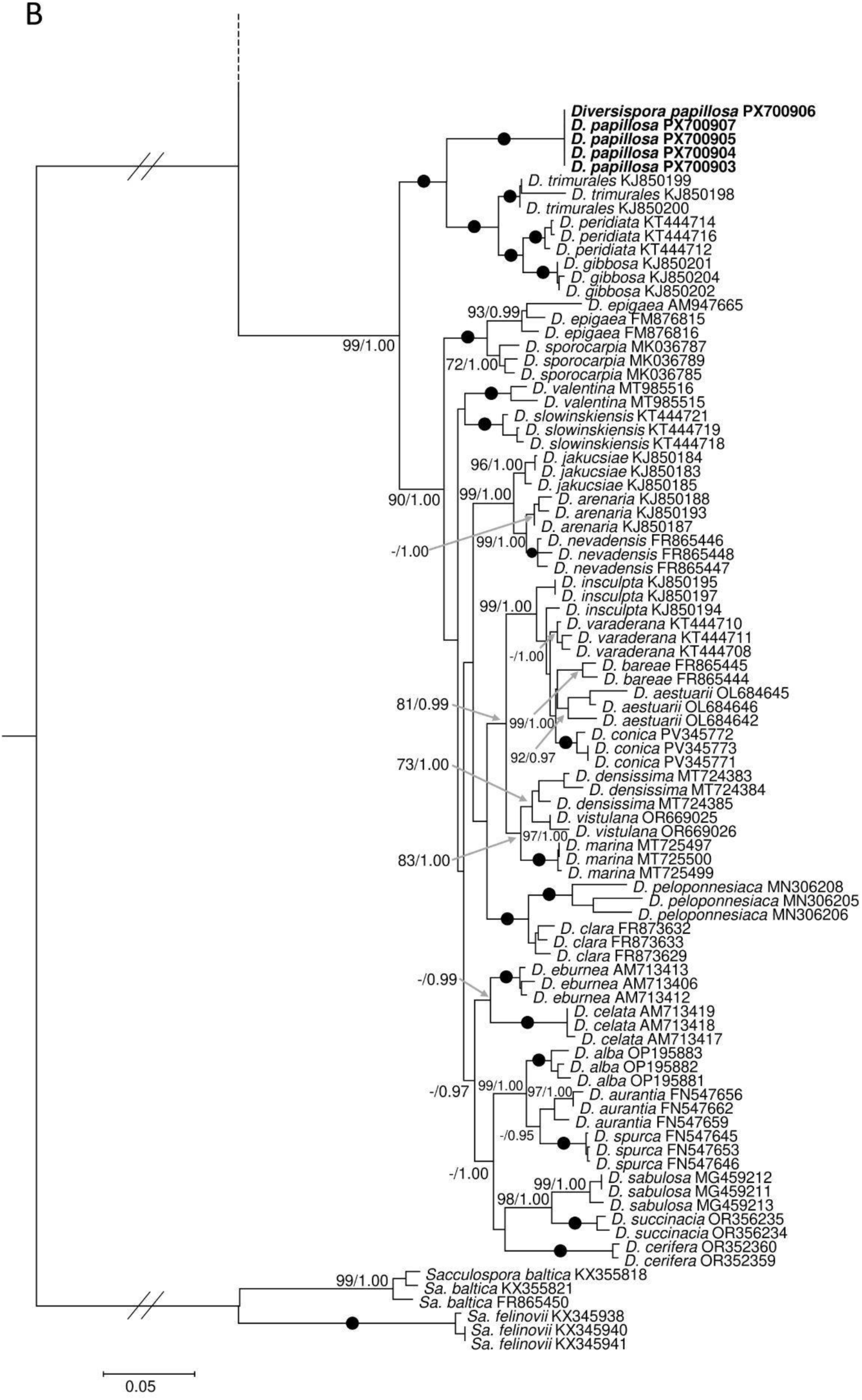
Phylogram generated from Maximum Likelihood (ML) and Bayesian Inference (BI) analyses displaying the phylogenetic relationships of the five new species (a,b), and the new genera and combinations (a) presented in the study. The new genera clades are displayed with colored boxes, while the new species and the sequences obtained in this study are highlighted in bold. *Sacculospora* species were used to root the tree. Posterior probabilities and support values ≥ 70% and 0.95, respectively, are indicated above or below the branches, while solid dots (●) on the branches represent full support (100/1.00). In panel b, all branches were equally stretched, and basal branches were shortened to 50% in length (indicated by //) to improve the visualization. In both panels, the bar indicates an expected change of 0.05 per site per branch.

Both phylogenies showed that the five specimens represented independent, fully or highly supported lineages within the order.

Furthermore, the three *Redeckera* reference species (*R. megalocarpa*, *R. fulva*, and *R. pulvinata*) split into three independent fully or highly supported lineages within the Corymbiglomeraceae. Given the polyphyly of *Redeckera sensu* C. Walker and A. Schüßler, the three clades are now recognized at genus rank, with *R. megalocarpa* clade retaining the name *Redeckera*, *R. fulva* clade becoming *Albocarpum,* and *R. pulvinata* clade as *Pulvinocarpum*. Furthermore, a fourth genus-rank lineage, hereafter referred to as *Melanocarpum*, was represented by sequences of the specimen “De”, hereafter referred to as *Melanocarpum mexicanum*. The four genera, together with *Corymbiglomus*, *Paracorymbiglomus,* and *Sieverdingia*, grouped into a supported clade (BI=1.00; ML=74), sister to *Desertispora*. Within this assemblage, none of the seven genera showed any supported sister relationship. The family Corymbiglomeraceae received full BI, and moderate ML support. The specimen “R2”, hereafter *Redeckera varelae*, was placed within the *Redeckera* clade, sharing 96-95.3% sequence identity with *R. megalocarpa*. Both “Fc2” and “Dt” specimens, hereafter *Albocarpum arenaceum* and *A. leptohyphum*, respectively, were placed in the *Albocarpum* clade, with *A. arenaceum* as the sister species of *A. fulvum*.

Finally, sequences of “Fc1”, hereafter *Diversispora papillosa*, formed a clade sister to *Diversispora trimurales*, *D. gibbosa,* and *D. peridiata*. Altogether, these species grouped into a fully supported clade, highly divergent (>7%) from the highly supported (BI=1.00; ML=90) sister clade hosting all the other *Diversispora* species.

In Supplementary Table 1, the intraspecific variability of the newly described species, as well as the intrageneric variability for all genera in Corymbiglomeraceae, is reported as percentages of sequence dissimilarity. Additionally, for both species and genera, the percentage of dissimilarity from the closest neighbour is provided.

The broader phylogeny (Figure 2), based on isolate-derived and eDNA sequences, supported the split of *Redeckera sensu* C. Walker and A. Schüßler into three independent genera, and confirmed the recognition of the new genus *Melanocarpum* along with the five new species described in this study. All four genera were fully supported, and, unlike in the phylogeny based on isolate-derived sequences, *Corymbiglomus* and *Pulvinocarpum* formed a fully supported sister relationship. These four genera, together with *Corymbiglomus* and *Paracorymbiglomus*, formed a highly supported clade (99) sister to *Sieverdingia*.

**Figure 2.**
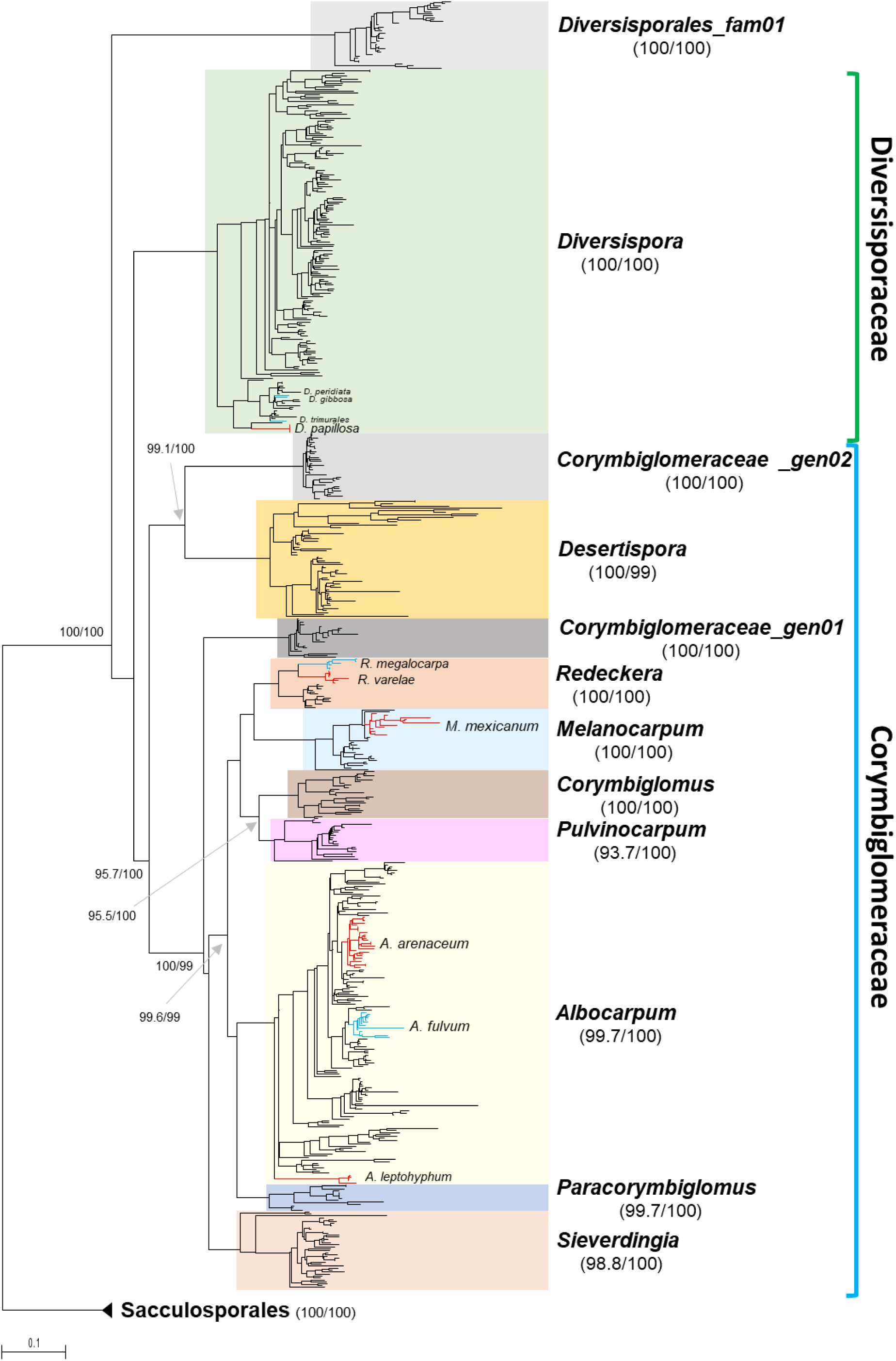
Phylogram generated from Maximum Likelihood analysis based on the eDNA dataset of Diversisporales and Sacculosporales (as out-group). The analysis conducted in IQ-TREE2 involved 685 sequences, overlapping the 45S barcode. Colored boxes highlight lineages at genus rank and the new candidate family in Diversisporales. The new species described in the paper are shown as red clades stemming from the most recent common ancestor (MRCA) to the sequences obtained in this study. Similarly, their close neighbours are shown as blue clades or branches. Genus-level support values (SH-aLRT and Ultrafast Bootstrap support) are shown beside the respective lineage labels, whereas support values for deeper nodes are indicated beside the branches. Bar indicates 0.1 expected change per site per branch.

In addition, one candidate new family and two novel genus-rank lineages, without representative isolates, were detected and provisionally designated as Diversisporales fam01, Corymbiglomeraceae gen01, and Corymbiglomeraceae gen02 (sister of *Desertispora*).

In the eDNA phylogeny, the newly presented species formed autonomous clades (Figure 2), most of which also included environmental sequences. Furthermore, the majority of eDNA sequences clustered into numerous additional clades, representing potentially new species, particularly in the *Albocarpum* lineage.

## Taxonomy

### Emendation of *Redeckera*

***Redeckera*** C. Walker & A. Schüßler emend. Magurno, Polo-Marcial & B.T. Goto

**MycoBank No:** xxx (will be added after reviews).

**Type species:** *Redeckera megalocarpa* (D. Redecker) C. Walker & A. Schüßler. The Glomeromycota, A species list with new families and new genera (Gloucester): 44. 2010.

**Basionym:** *Glomus megalocarpum* D. Redecker Mycol. Progr. 6: 38. 2007. **Diagnosis:** It differs from other genera in the Corymbiglomeraceae in (i) the formation of large, irregularly compact or pulvinate glomerocarps (> 5 mm diam.), with defined peridium and gleba, cottony in appearance and containing thousands of spores, (ii) glomerospores ovoid with a one-to three-layered wall, (iii) subtending hypha (SH) straight or slightly curved, concolorous with the spore wall, wide at the base and narrowing as they extend, formed by 1 or 2 layers, (iv) SH occluded by a thick, straight septum at or up to 2 μm below the spore base, and (v) the nucleotide composition of 45S sequences.

**Genus description:** See diagnosis (above) and Oehl et al. (2011a) for *Redeckera*.

**Distribution and habitat:** In the field, environmental sequences from EUKARYOME v.1.9.4 suggest that, although the number of available records is not high, they originate from tropical, subtropical, and temperate regions. Tropical records originate from Puerto Rico and the Dominican Republic, all associated with tropical broadleaf forests. In temperate regions, the only available records are from Estonia, where the genus occurs predominantly in woodlands. There are also records from Mexico, associated with subtropical forests, including both coniferous and broadleaf formations (Supplementary Figure 1A).

### Description of a new species

*Redeckera varelae* Polo-Marcial, Magurno & B.T. Goto, sp. nov. Figure 3 A–H.

**Figure 3.**
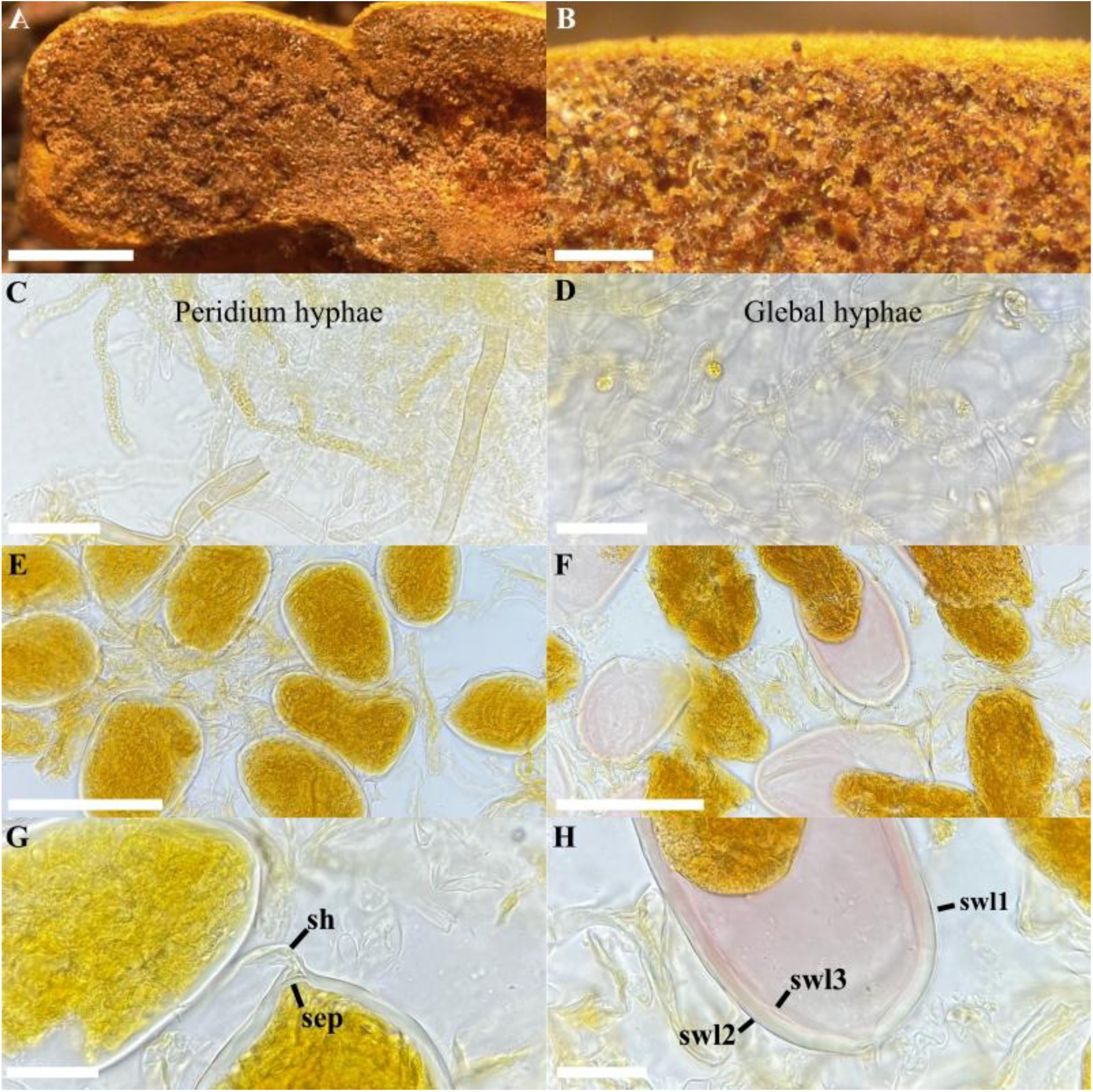
*Redeckera varelae*. (A, B) fresh glomerocarp with glomerospores, (C) detail of the peridium hyphae, (D) detail of the glebal hyphae, (E) intact glomerospores, (F) detail of the granulated cytoplasmic content, (G) subtending hypha (sh) and septum (sep), (H) spore wall layers (swl)1 – 3. (D, E, G) hyphae and glomerospores in PVLG, (C, F, H) hyphae and glomerospores in PVLG+Melzer’s reagent. Scale bars: (A) = 2 mm, (B) = 300 μm, (C, D, G, H) = 20 μm, (E, F) = 100 μm

**MycoBank No:** xxx (will be added after reviews).

**Etymology**: Latin, *varelae*, in honor of Dr. Lucía Varela Fregoso in recognition of her important contributions to the taxonomy of Glomeromycota in Mexico.

**Specimens examined**: Chetumal, Quintana Roo, Mexico. Glomerocarp isolated from a highly disturbed area within the Zazil Ecological Urban Park by Javier de la Fuente and Hassan Polo on October 27, 2023. Holotype: (will be inserted after review), isotype (will be inserted after review) and paratype: (will be inserted after review).

**Diagnosis**: Differs from *R. megalocarpa*, the other species in the genus, by (i) glomerospores with a three-layered wall, (ii) the second layer staining pink in Melzer’s reagent, (iii) dense, granular yellow cytoplasmic content, and (iv) the nucleotide composition of sequences of the 45S marker.

**Description**: Glomerocarps sub-hypogeous, compact and pulvinate, with delimited peridium and gleba. Peridium cottony, pale yellow (4A7) to mustard yellow (4B8), 9–14 × 4–5 mm (Figure 3A, B), with interwoven hyphae, sub-hyaline to light yellow (1A2), 2.0–5.4 μm wide, with a 0.5–0.8 μm thick wall (Figure 3C). Gleba light yellow (1A2) to golden (2A4), with straight or branched dark-pigmented hyphae 5.0–7.4 μm wide, and wall with 0.7–1.4 μm thickness (Figure 3D). Glomerospores light brown to pale brown (3A8), due to the cytoplasmic contents densely granulated, developing blastically at the tip of a subtending hypha, usually ovoid, 71–100 × 40–67 μm, rarely subglobose, (62–)74(–80) μm diam., or clavate, 90–120 × 49–62 μm diam. (Figure 3E, F). Glomerospores with three wall layers (SWL1–3) (Figure 3H). Layer 1 uniform (no sublayers visible), semi-flexible, smooth, permanent and hyaline, 0.5–0.8 μm thick (Figure 3H), strongly adherent to SWL2. Layer 2 permanent, laminated, hyaline to light yellow (1A2), (2.6–)3.8(–5.1) μm thick (Figure 3H). Layer 3 hyaline, permanent, semi-flexible, and strongly adherent to SWL2, 0.5–0.7 μm thick. Only layer 2 stains pink (14A3) in Melzer’s reagent (Figure 3F, H). Subtending hypha fragile, concolorous and continuous with spore wall layers, usually detached upon mounting; (25–)36(–45) μm long, straight or curved, cylindrical or funnel-shaped (Figure 3G). Wall of subtending hypha formed by two layers (SHWL1–2) concolorous with the spore wall; SHWL1 0.5–0.8 μm thick, deteriorated or completely sloughed off in mature spores; SHWL2 1.0–1.5 μm thick (Figure 3G). Pore (5.4–)7(–11) μm wide at the spore base, occluded by a septum (1.0–)1.5(–2.3) thick, formed by the SWL2–3 (Figure 3G, H). The position of the septum is up to 2.5 μm above the termination of SWL2 (Figure 3G, H). Spore content of light brown to pale brown (3A8) oily substance. Germination unknown.

**Distribution and habitat**: In the field, *R. varelae* has been detected only in preserved and disturbed fragments of tropical semi-evergreen forest, on the soil surface or attached to organic debris and on primary roots of *Lysiloma latisiliquum* (L.) Benth. (de la Fuente et al. 2023). BLASTn search in GenBank did not return any potential match. In EUKARYOME v.1.9.4, two ITS sequences with a high percentage of identity (96.5-100%) extended the known distribution of *R. varelae* to tropical broadleaf forests in Puerto Rico (EUK0089452) and the Dominican Republic (EUK0510416).

## Other species

*Redeckera megalocarpa* (D. Redecker) C. Walker & A. Schüßler, The Glomeromycota: a species list with new families and new genera: 44 (2010).

≡ *Glomus megalocarpum* D. Redecker, Mycol. Progr. 6 (1): 38 (2007).

*Redeckera avelingiae* (R.C. Sinclair) Oehl, G.A. Silva & Sieverd., Mycotaxon 116: 111 (2011).

≡ *Glomus avelingiae* R.C. Sinclair, Mycotaxon 74: 338. 2000.

*Redeckera canadensis* (Thaxt.) Oehl, G.A. Silva & Sieverd., Mycotaxon 116: 111 (2011).

≡ *Endogone canadensis* Thaxt., Proc. Am. Acad. Arts Sci. 57: 317. 1922.

≡ *Glomus canadense* (Thaxt.) Trappe & Gerd., Mycol. Mem. 5: 59. 1974.

*Redeckera fragilis* (Berk. & Broome) Oehl, G.A. Silva & Sieverd., Mycotaxon 116: 111 (2011).

≡ *Paurocotylis fragilis* Berk. & Broome, J. Linn. Soc. Bot. 14: 137. 1873.

≡ *Glomus fragile* (Berk. & Broome) Trappe & Gerd., Mycol. Mem. 5: 59. 1974.

**Erection of a new genus, species and new combinations *Albocarpum*** Magurno, Polo-Marcial, Crossay & B.T. Goto gen. nov. **MycoBank No:** xxx (will be added after reviews).

**Etymology:** Latin, *albo* (= white) and *carpum* (= fruitbody), in reference to the color of the glomerocarps (= sporocarps).

**Type species:** *Albocarpum leptohyphum* Polo-Marcial, Magurno, B.T. Goto sp. nov. **Diagnosis:** It differs from other genera in the Corymbiglomeraceae in (i) production of glomerospores in compact glomerocarps with delimited peridium and gleba enclosing thousands of spores and in loose clusters of up to 15 spores, (ii) globose to subglobose, ellipsoid or ovoid glomerospores with a three-layered wall, (iii) phenotypic and histochemical composition of the wall, (iv) a straight, cylindrical or funnel-shaped subtending hypha composed of one or two layers and occluded by a straight septum at the spore base, and (v) the nucleotide composition of sequences of 45S barcode.

**Genus description:** Glomerospores globose to subglobose, (70‒)90(‒114) μm diam, produced in compact glomerocarps, hypogeous, and pulvinate 6–11 × 0.7–1.1 μm thick with cottony peridium formed by intertwined hyphae, and containing thousands of randomly formed glomoid spores or in clusters of 5 to 15 spores in soil. Subtending hypha hyaline to white, fragile, straight or rarely recurved, cylindrical or slightly funnel-shaped, formed by one or two layers and occluded by a straight septum.

**Distribution and habitat:** In the field, according to the EUKARYOME v.1.9.4 database, the genus is broadly distributed along the tropical belt and its closely subtropical adjacent zones, with records spanning multiple continents and a wide range of biomes. In Africa, it occurs in broadleaf forests (South Africa, Madagascar, Benin, Ghana, Gabon, Rwanda), tropical woodlands (Benin, Ghana, Senegal, Tanzania), tropical coniferous forests (Ethiopia), tropical grasslands (Gabon), and croplands (Côte d’Ivoire). In Asia, it has been reported from tropical and subtropical broadleaf forests (India, Thailand, Indonesia, Philippines, Saudi Arabia), tropical grasslands (Bangladesh), croplands (Thailand), and desert habitats (Saudi Arabia). In Oceania, records come from tropical broadleaf forests and croplands in Papua New Guinea, tropical broadleaf forests in Australia, and from tropical shrubby vegetation and cropland in the Pacific atoll of French Polynesia. In the Americas and Caribbean, it is found mainly in broadleaf forests across Brazil, Colombia, Mexico, Panama, Costa Rica, Argentina, Cuba, USA, Puerto Rico, Guadeloupe, Dominica, and the British Virgin Islands (Supplementary Figure 1B).

### Description of new species

*Albocarpum leptohyphum* Polo-Marcial, Magurno & B.T. Goto, sp. nov. Figure 4A‒F.

**Figure 4.**
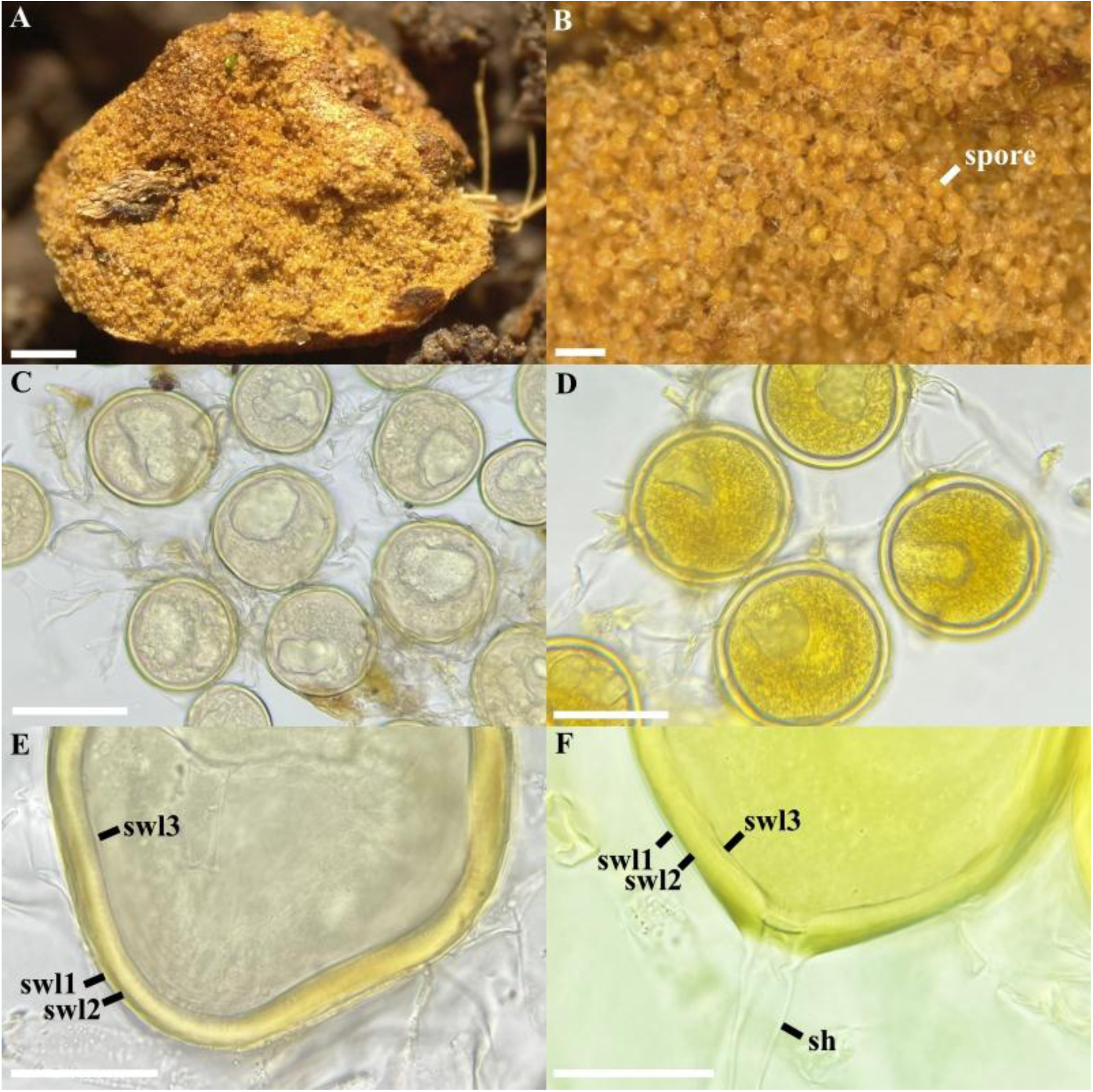
*Albocarpum leptohyphum*. (A) Lateral view of the glomerocarp, (B) glomerospores formed between the gleba, (C, D) intact glomerospores with subtending hypha, (E, F) Spore wall layers (swl) 1 – 3, and subtending hypha (sh). (C, E) glomerospores in PVLG, (D, F) glomerospores in PVLG+Melzer’s reagent. Scale bars: (A) = 1 mm, (B) = 200 μm, (C) = 100 μm, (D) = 50 μm, (E, F) = 20 μm.

**MycoBank No:** xxx (will be added after reviews).

**Etymology:** Greek, λεπτός, *leptós* (= delicate) and Latin, *hyphum* (= hypha), in reference to the fragility of the subtending hypha.

**Specimens examined**: Valle del Ovni, Gómez Farías, Tamaulipas, Mexico. Epigeous glomerocarps, solitary, attached to root fragments in tropical mountain cloud forest (23°01’26.2”N, 99°09’57.5”W, 540 masl) by Javier de la Fuente on July 20, 2024.

Holotype: (will be inserted after review), isotype (UFRN-FUNGOS, will be inserted after review).

**Diagnosis:** It differs from other species of the genus *Albocarpum* in (i) the formation of globose to subglobose, light yellow to golden yellow spores, (ii) the reaction of the second layer and the cytoplasmic content to Melzer’s reagent, and (iii) the nucleotide composition of 45S sequences.

**Description:** Glomerocarps compact, hypogeous, and pulvinate; light brown (5C5) to pale brown (5E5), 4–5 × 2.5–3.5 mm, with a thin, cottony peridium formed by intertwined hyphae, pale yellow (3A7) to light brown (4B7), 6–11 μm wide, 0.7–1.1 μm thick (Figure 4A) and containing thousands of randomly formed glomoid spores (Figure 4B‒C). Sub-hyaline to light yellow (1A6) gleba formed by intertwined hyphae 4–11 μm wide and with a wall 1–1.2 μm thick (Figure 4B, C). Globose to subglobose glomerospores, (70‒)90(‒114) μm diam., light yellow (1A6) to golden yellow (2A5) (Figure 4C, D). The spore wall consists of three layers (Figure 4E, F). The first layer (SWL1) subhyaline, unitary, permanent, smooth, and semi-flexible, 0.5–1.3 μm thick. The second layer (SWL2) laminated, golden yellow (2A5), 4–8 μm thick. The third layer (SWL3) permanent and semi-flexible, 1–1.5 μm thick, concolored and strongly adhered to SWL2 (Figure 4E, F). In Melzer’s reagent, the SWL2 and the cytoplasmic content stain intense yellow (2A8). Subtending hypha hyaline to white (1A1), fragile, and often detached into spores when pressure is applied; straight or rarely recurved and cylindrical, 8–10 μm wide at the spore base, formed by two hyaline layers continuous with SWL1‒2 (Figure 4E, F). The SHWL1 1–1.2 μm thick and SHWL2 1–1.5 μm thick. Pore (6‒)8(‒11) μm wide, occluded by a straight, thick septum (1–1.5 μm thick), formed by the SWL2 and SWL3. The septum is often located halfway through the thickness of the SWL2 (Figure 4E, F). Spore content of hyaline to light yellow oily substance (1A3). Germination unknown.

**Distribution and habitat:** In the field, *A. leptohyphum* has only been reported from the type locality in the tropical mountain cloud forest of Northern Mexico. Neither BLASTn analysis in GenBank nor in EUKARYOME v.1.9.4 returned any match.

*Albocarpum arenaceum* T. Crossay, M. Wong, Magurno & B.T. Goto, sp. nov. Figure 5 A‒F.

**Figure 5.**
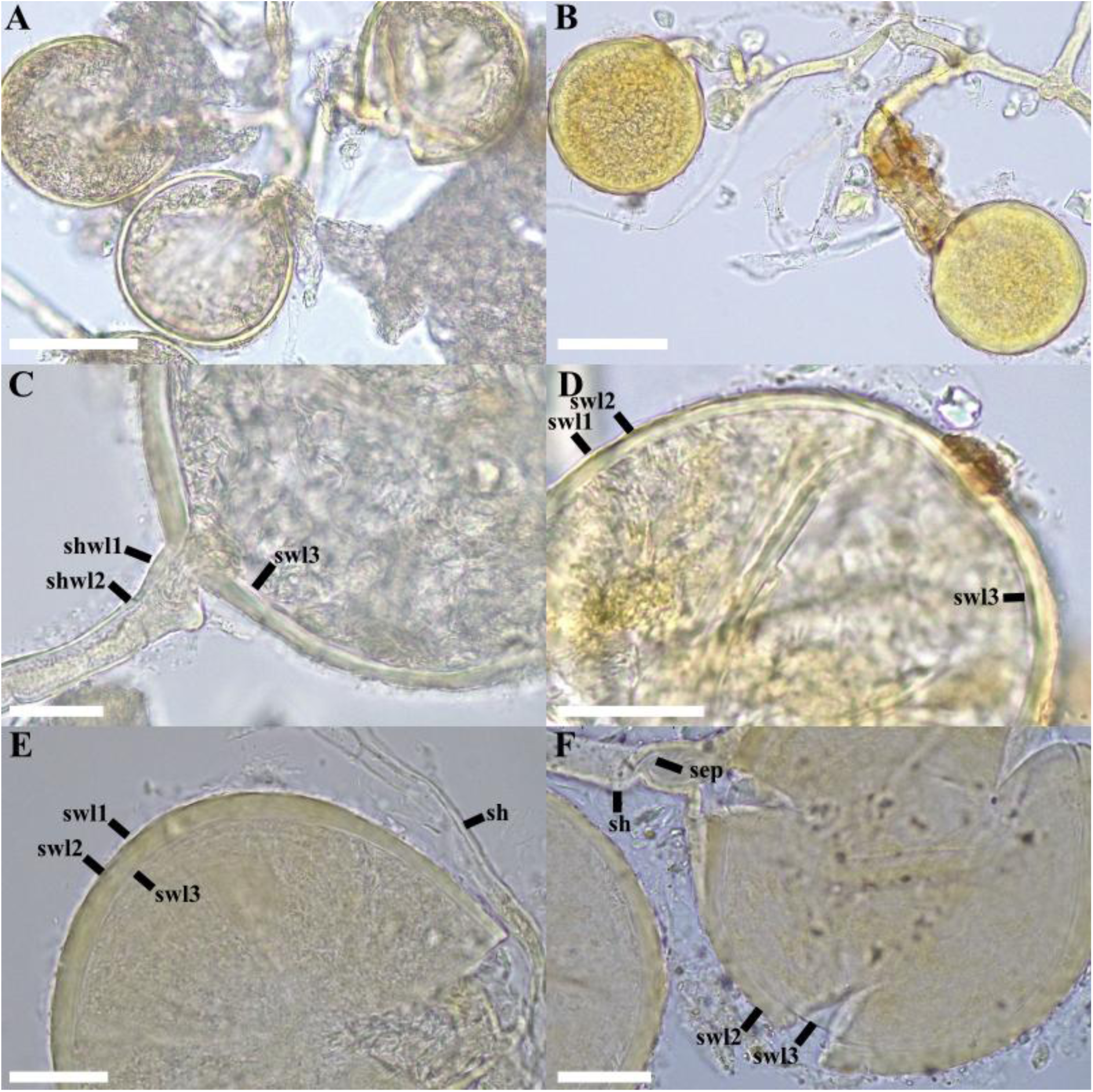
*Albocarpum arenaceum*. (A, B) Glomerospores produced in loose clusters. (C-F) Spore wall layers (swl) 1 – 3. (E,F) Subtending hypha (sh) and septum (sep). (A, B, C, E, F) in PVLG; (D) in PVLG + Melzer’s reagent. Scale bars: (A, B) = 50 μm, (C, D) = 25 μm, (E, F) = 20 μm.

**MycoBank No:** xxx (will be added after reviews).

**Etymology:** Latin, *arenaceum* (= sandy), in reference to the “sandy” spore appearance.

**Specimens examined**: Isolated from rhizospheric soil of a greenhouse pot of a single–species culture propagated on *Sorghum bicolor* (L.) Moench at the laboratory Aura Pacifica in New Caledonia, Noumea, June 2024, T. Crossay. The single–species culture was originally inoculated with 100 spores isolated from the rhizospheric soil of *Ficus carica* L. sampled at a naturally vegetated site located on Rangiroa Island (14°58’24.3“S, 147°37’28.4”W) in French Polynesia, October 2022, T. Crossay. Holotype deposited at the Mycological Herbarium of the National Museum (France, Paris) in MNHN–PC–(will be inserted after review), isotype deposited (will be inserted after review).

**Diagnosis:** It differs from other species of the genus *Albocarpum* in (i) the formation of spores in loose aggregates, (ii) the phenotypic and histochemical properties of the wall, (iii) the funnel-shaped subtending hypha, and (iv) the nucleotide composition of the 45S sequences.

**Description:** Glomerocarps unknown. Glomerospores produced in clusters of 5 to 15 spores in soil, arising at the tips of sporogenous hyphae branched from a parent hypha continuous with extraradical hyphae or sporogenous hyphae directly continuous with extraradical hyphae (Figure 5A, B). Spores globose to subglobose, off-white (4A2) to light buttercup yellow (4A7), (80–)90(–120) μm diam., with one subtending hypha (Figure 5A, B, C, E, F). Spore wall composed of three layers (Figure 5D, E, F). Layer 1 (SWL1) forming the spore surface, evanescent, mucilaginous, hyaline, (1.0–)1.2(–1.4) μm thick (Figure 5C, D, E, F), completely sloughed off in mature spores. Layer 2 (SWL2) uniform, permanent, smooth, semi-flexible, laminate, light yellow, (1A5), (4.0–)5.4(–7.0) μm thick (Figure 5C-F). Layer 3 (SWL3) flexible, hyaline and difficult to detect even if spores break under hard pressure (Figure 5C-), (1.5–)2.5(–3.5) μm thick. Layers 1–3 without Melzer’s reaction (Figure 5B, E). Subtending hypha hyaline to light yellow (1A5), straight, cylindrical, rarely slightly constricted at the spore base. The subtending hyphal wall consists of three layers continuous with the spore wall layers, (2.0–)2.5(–3.0) μm thick at the base of the spore, extending to the distal end of the hypha (Fig. C, F). The pore has a diameter of (4.0–)4.5(–5.0) μm, occluded by a curved septum, 1–1.3 μm thick, formed by SWL3 and extending to 5–8 μm below the base of the spore (Figure 5E, F). Spore content of off-white (4A2) to yellow (4A7), dense oily substance. Germination unknown. Forming hyphal coils and arbuscules, staining with Trypan Blue in the root cortex of *Sorghum bicolor*.

**Distribution and habitat:** In the field, *A. arenaceum* has been reported only from its type locality on Rangiroa Island, French Polynesia. BLASTn searches in GenBank returned several sequences with 97-98% identity, from an AMF community associated with *Hevea brasiliensis* (Willd. ex A.Juss.) Müll.Arg. in Kerala (e.g., PQ669996, PQ669944; India). In EUKARYOME v.1.9.4, many potential occurrences of the species were detected. Considering BLASTn matches with the highest percentage of identity (97-98.6%), *A. arenaceum* could be found in tropical-subtropical forests (Colombia, EUK1675185; British Virgin Islands, EUK1675202; Benin, EUK1675200; South Africa, EUK1675134; Gabon, EUK1675138; and India, EUK1675143) and subtropical deserts (Saudi Arabia, EUK1675213).

### New combination

*Albocarpum fulvum* (Berk. & Broome) Magurno, Polo-Marcial & B.T. Goto, comb. nov. Figure 6 A-F.

**Figure 6.**
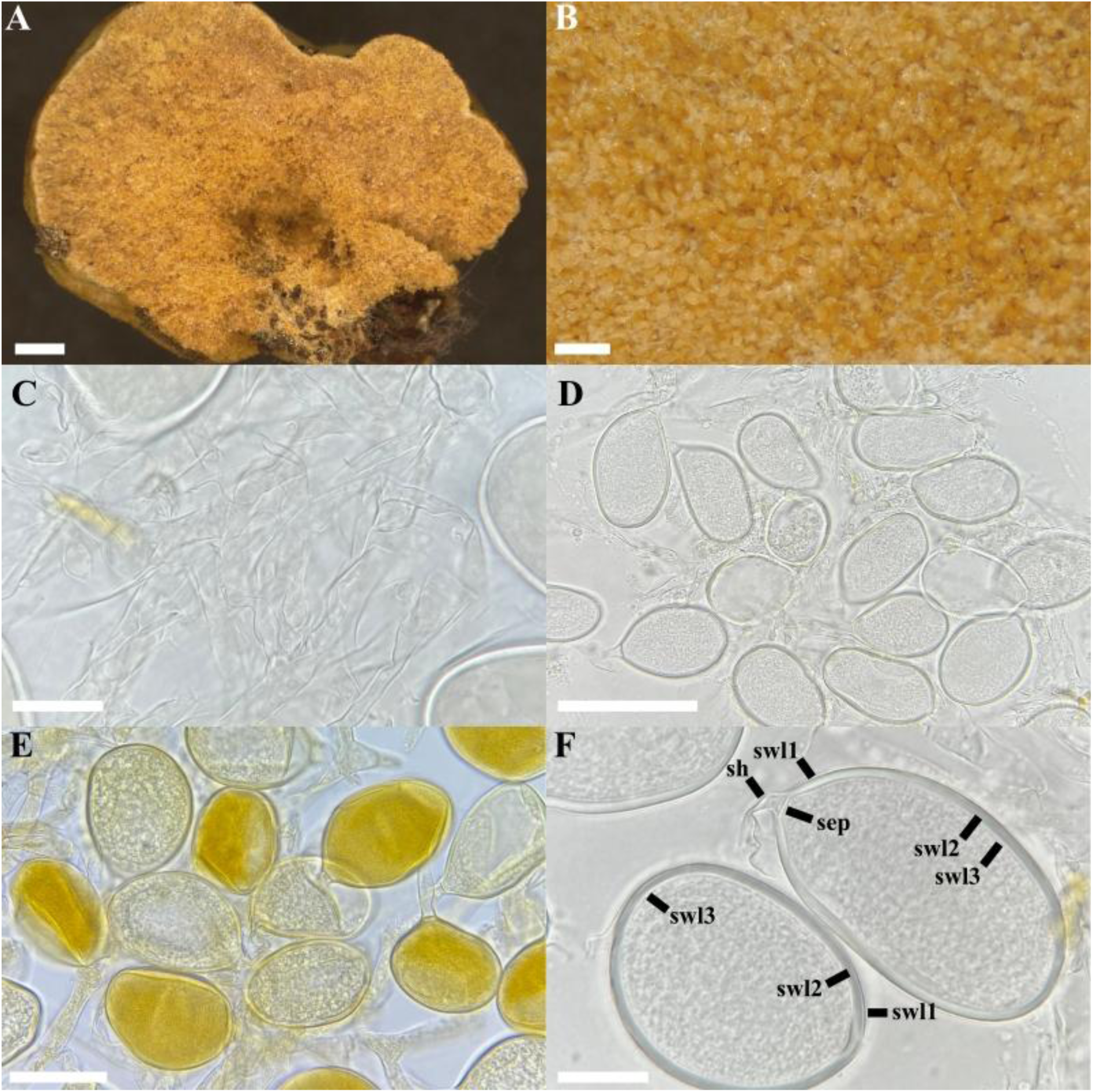
*Albocarpum fulvum*. (A) Lateral view of the glomerocarp, (B) glomerospores formed in the gleba, (C) hyphae of the gleba, (D) intact glomerospores in PVLG, (E) intact glomerospores in PVLG + Melzer, (F) spore wall layers (swl) 1 – 3, and subtending hypha (sh) with septum (sep). Scale bars: (A) = 1 mm, (B) = 200 μm, (D) = 100 μm, (C, E, F) = 20 μm.

**MycoBank No:** xxx (will be added after reviews).

**Specimens examined**: Petcacab, Quintana Roo, Mexico. Solitary or gregarious glomerocarps attached to roots and leaf litter fragments in a preserved fragment of evergreen tropical forest (19°12’02.5“N, 88°26’54.0”W, 30 masl) by Hassan Polo on October 28, 2023. Holotype: (will be inserted after review), isotype (will be inserted after review).

**Basionym:** *Redeckera fulva* (Berk. & Broome) C. Walker & A. Schüßler, The Glomeromycota, A species list with new families and new genera (Gloucester): 44. 2010.

≡ *Glomus fulvum* (Berk. & Broome) Trappe & Gerd., Mycol. Mem. 5: 59. 1974.

≡ *Paurocotylis fulva* Berk. & Broome, J. Linn. Soc. Bot. 14: 137. 1873.

**Diagnosis**: It differs from other *Albocarpum* species in (i) the formation of frequently ellipsoid spores, (ii) a three-layered wall, whose laminated layer (SWL2) is pigmented light yellow (1A2) in Melzer’s reagent, (iii) a subtending hypha with a wide channel occluded by a straight septum formed at the base of the spore, and (iv) the nucleotide composition of the partial 45S sequences.

**Epitype description**: Glomerocarps compact, irregular to pulvinate, white (1A1) with a cottony texture when fresh, hours later they acquire a beige (5A2) to pale brown (5D7) color, 2–9 × 4–15 mm (Figure 6A). Peridium sub-hyaline to golden yellow (1A7) on Melzer, formed by randomly intertwined hyphae 3–4.5 μm wide and 0.5–0.8 μm wall thickness (Figure 6B). Glomerospores are randomly arranged in the gleba with hyphae 8–15 μm wide and 0.8–1.0 μm wall thickness. Hyaline to light yellow (1A2), concolorous with subtending hyphae (Figure 6B, D, F), frequently ellipsoid (66–)70 × 90(–113) μm, rarely ovoid (62–)69 × 84(–88) μm or subglobose 76 × 75 μm diam. Spore wall consists of three layers: SWL1 hyaline to light yellow, 0.5–1.0 μm thick, permanent and smooth (Figure 6F); SWL2 concolorous and strongly adhered to SWL1, laminated, (1.5–)2.5(–3) μm thick; SWL3 concolorous, strongly adhered to SWL2, 0.7–1.3 μm thick, permanent and smooth (Figure 6F). Subtending hypha concolorous and continuous with SWL1–2, (6–)15(–27) μm length, straight or curved, cylindrical or funnel-shaped (4–)5(–8) μm wide at base of spore. SHWL1 is rarely observed; when present, it is very thin, 0.5 μm thick; SHWL2 0.8–1 μm thick, occluded by a septum formed by SWL2 (Figure 6F). Spore content of hyaline oily substance.

Spore wall and cytoplasmic contents acquire a slight yellow (1A2) coloration in Melzer’s reagent (Figure 6E).

**Distribution and habitat**: In the field, environmental sequences from EUKARYOME v.1.9.4 with 97-99% identity indicate several possible occurrences of the species in Caribbean tropical forests (Mexico, EUK1675191; Costa Rica, EUK0512176; Panama, EUK1675251; Guadalupe, EUK1675187; Puerto Rico, EUK0512172; and Colombia, EUK1675183).

**New combination and erection of a new genus *Pulvinocarpum*** Polo-Marcial, Magurno & B.T. Goto gen. nov. **MycoBank No:** xxx (will be added after reviews).

**Etymology:** Latin, *pulvino* (= pillow) and *carpum* (= fruitbody), in reference to the pillow-shaped glomerocarps (= sporocarps).

**Type species:** *Pulvinocarpum pulvinatum* (Henn.) B.T. Goto, Polo-Marcial & Magurno comb. nov.

**Basionym:** *Redeckera pulvinata* (Henn.) C. Walker & A. Schüßler, The Glomeromycota: 44. 2010.

≡ *Glomus pulvinatum* (Henn.) Trappe & Gerd., Mycol. Mem. 5: 59. 1974.

≡ *Endogone pulvinata* Henn., Hedwigia 36: 212. 1897.

**Diagnosis:** It differs from other genera in the Corymbiglomeraceae as (i) it produces compact glomerocarps, with a small peridium, containing hundreds of disorganized spores, (ii) glomerospores are globose or subglobose, rarely ellipsoid, composed of a three-layered wall with granular cytoplasmic material, and without Melzer’s reaction, (iii) a wide subtending hypha at the spore base, concolorous and continuous with the first and second layers of the spore wall, occluded by a thick, straight septum formed by the innermost layers, (iv) the layers commonly form folds in the center of the spore, even without applying pressure, (v) in addition to its nucleotide composition of sequences of the 18S-ITS nuc rDNA region.

**Genus description:** See diagnosis (above).

**Distribution and habitat:** In the field, environmental sequences indicate that the genus is widely distributed across tropical and subtropical regions, occurring in both natural and anthropogenic habitats. Records are from tropical broadleaf forests in New Caledonia, Dominica, and Colombia; subtropical broadleaf and tropical coniferous forests and urban environments in Mexico; subtropical grasslands in Uruguay; subtropical shrublands in the Canary Islands; and tropical woodlands and xeric shrublands in Brazil (Supplementary Figure 1C).

### Erection of a new genus and species

***Melanocarpum*** B.T. Goto, Polo-Marcial, M.B. Queiroz & Magurno gen. nov.

**MycoBank No:** xxx (will be added after reviews).

**Etymology**: Latin, *melano* (= dark-coloured) and *carpum* (= fruitbody), in reference to the color of the glomerocarps (= sporocarps).

**Type species**: *Melanocarpum mexicanum* Polo-Marcial, Magurno & B.T. Goto, sp. nov.

**Diagnosis**: It differs from other genera of Corymbiglomeraceae in the (i) production of glomerospores in compact and disorganized glomerocarps, (ii) large globose glomerospores (250 μm), (iii) phenotypic characteristics of the spore wall, (iv) Melzer reaction, and (v) the nucleotide composition of 45S sequences.

**Genus description**: Producing glomoid spores in compact, semi-hypogeous glomerocarps with a thin peridium. Golden yellow to light orange spores, globose to subglobose, 180–290 µm diam., with a four-layered wall; only the second layer (laminated) stains dark orange in Melzer’s reagent. Subtending hypha cylindrical, slightly funnel-shaped, color discontinued to spore wall, with a wall composed of three continuous layers with the spore wall, except for the innermost layer. Subtending hyphal pore closed by a straight septum continuous with the innermost layer of the spore wall.

**Distribution and habitat:** In the field, EUKARYOME v.1.9.4 metadata indicate a broader distribution across Mexico and the Caribbean islands, including tropical and subtropical broadleaf forests in Cuba, Puerto Rico, the Dominican Republic, and Dominica (Supplementary Figure 1D).

*Melanocarpum mexicanum* Polo-Marcial, Magurno & B.T. Goto, sp. nov. Figure 7A–H.

**Figure 7.**
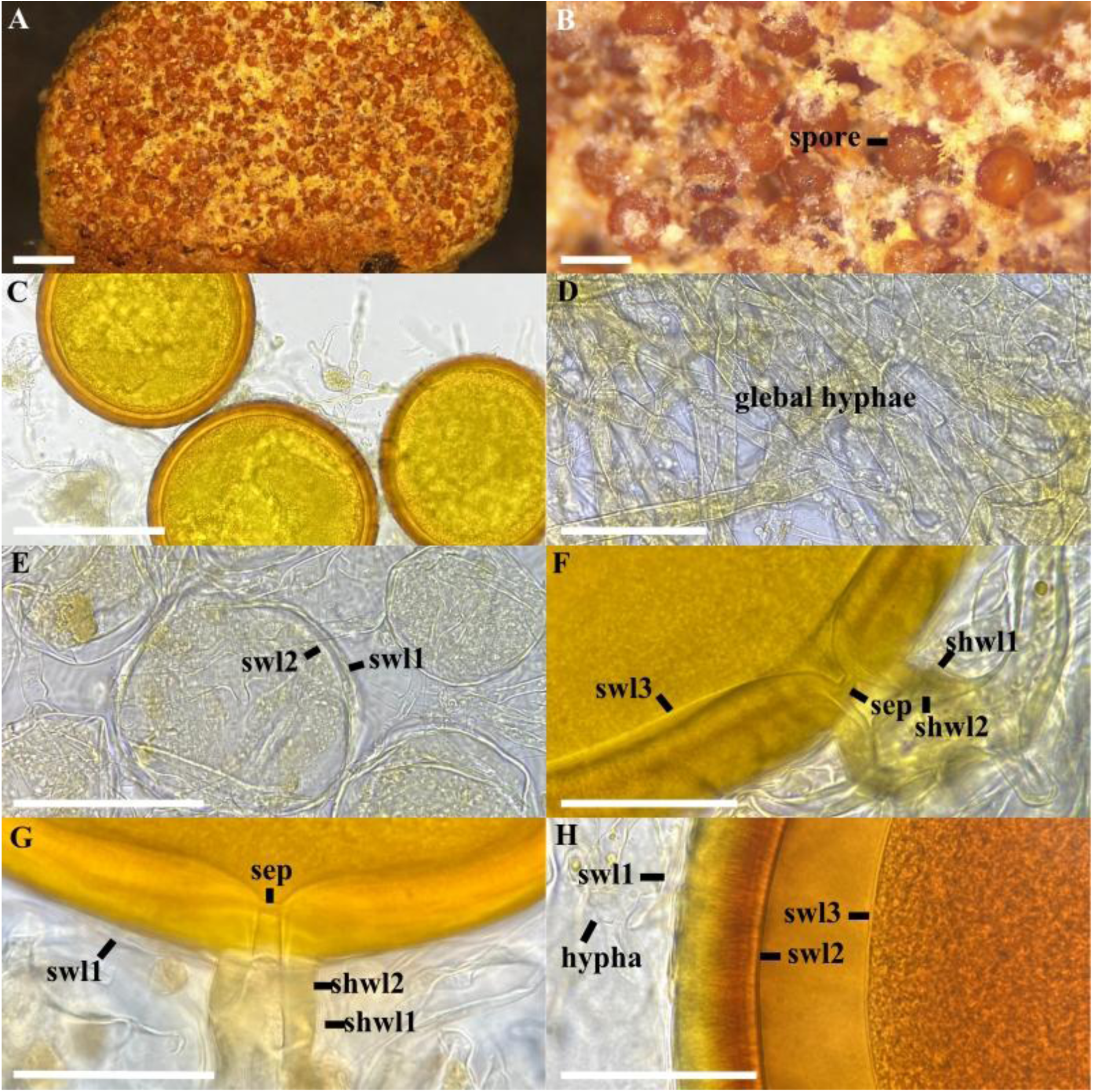
*Melanocarpum mexicanum*. (A, B) Glomerocarp with hundreds of spores formed between the gleba, (C) intact glomerospores, (D) detail of the glebal hyphae, (E) immature glomerospores with a two-layered wall (swl) 1 – 2, (F, G) subtending hypha with two wall layers (shwl) 1 – 2 and septum (sep). (H) spore wall layers (swl) 1 – 3. (C, E–G) glomerospores in PVLG, (D, H) glomerospores in PVLG+Melzer’s reagent. Scale bars: (A) = 1 mm, (B–F) = 200 μm, (F–H) = 50 μm.

**MycoBank No:** xxx (will be added after reviews).

**Etymology**: Latin, *mexicanum*, in reference to Mexico, the country where the new species was originally found.

**Specimens examined**: Galeana, Campeche, Mexico. Glomerocarps gregarious or coalescent in a fragment of tropical deciduous forest (18°10’42.7“N, 89°14’30.2”W) by Javier de la Fuente on October 01, 2023. Holotype: (will be inserted after review), isotype (will be inserted after review).

**Diagnosis**: As that regarding the genus *Melanocarpum* (see above).

**Description**: Glomerocarps compact, unorganized and semi-hypogeous. Glomerocarps with a thin, cottony-looking peridium, 8–15 × 4–7 mm (Figure 7A). Light orange (5A4) to light brown (5C7) peridium, absent over time, usually partially covering the glomerocarps; formed by intertwined hyphae 3–7 µm thick. Whitish (1A1) to pale yellow (4A5) gleba; formed by straight or branched hyaline to golden hyphae (2A4); 3.8–15 µm wide and 1–1.5 µm thick; contains hundreds of mature and abortive spores, subhyaline to golden yellow, smooth, without apparent laminations (Figure 7B, D and E). The peridium and gleba do not react in Melzer’s reagent. Glomerospores arising blastically at tips of subtending hypha; golden yellow (2A6) to orange (5A8); globose to subglobose (180–)250(–290) µm diam (Figure 7A-C). The wall consists of three layers (Figure 7B, C). SWL1 sub-hyaline to light yellow (2A4), semi-flexible, semi-permanent, and smooth, intact in young glomerospores, generally deteriorates and breaks down into mature spores, 1.5–2 µm thick when intact (Figure 7G). SWL2 laminated, with fibrillar appearance and opacity in mature spores, permanent, yellow (2A7) to light orange (4A6), (16–)20(–23) µm thick; and rough surface appearance under light pressure (Figure 7H). SWL3 permanent, flexible, hyaline, 0.5–1.5 µm thick, in PVLG it folds and appears to be a double layer (Figure 7H). In abortive spores, the thickness of the wall layers is 0.7–1 (SWL1) and 0.5–1 (SWL2) µm, respectively. In Melzer’s reagent, only layer 2 stains dark orange (6B8) (Figure 7H). Subtending hypha cylindrical, slightly funnel-shaped, sub-hyaline to light yellow (2A4), (18–)25(–30) μm wide at spore base (Figure 7F, G). Composed of three layers (SHWL), concolorous, sub-hyaline to light yellow (2A4). SHWL1 generally deteriorated or detached 0.5–1 µm thick; SHWL2 (4.5 –)6(–8.7) μm thick; SHWL3 strongly adhered to the previous layer, 0.5 µm thick. Pore (5–)10(–13) µm diam., occluded by a straight or rarely slightly curved septum formed by SHWL1–3 (Figure 7F, G) located up to 8.5 µm below the base of the spore, where SHWL3 is difficult to observe (Figure 7G). Spore content of hyaline to light yellow oily substance (1A4). Germination unknown.

**Distribution and habitat**: In the field, *M. mexicanum* has been found only at the type locality in the tropical deciduous forest of Campeche, Mexico. BLASTn search in GenBank did not return any potential match. In EUKARYOME v.1.9.4, sequences with a high percentage of identity (97-99%) were obtained from four different locations with subtropical forest vegetation, all in Mexico (EUK1675189, EUK1675175, EUK1675197, EUK0089432).

### Description of a new species

***Diversispora papillosa*** T. Crossay, M. Wong & B.T. Goto, sp. nov. Figure 8A–F.

**Figure 8.**
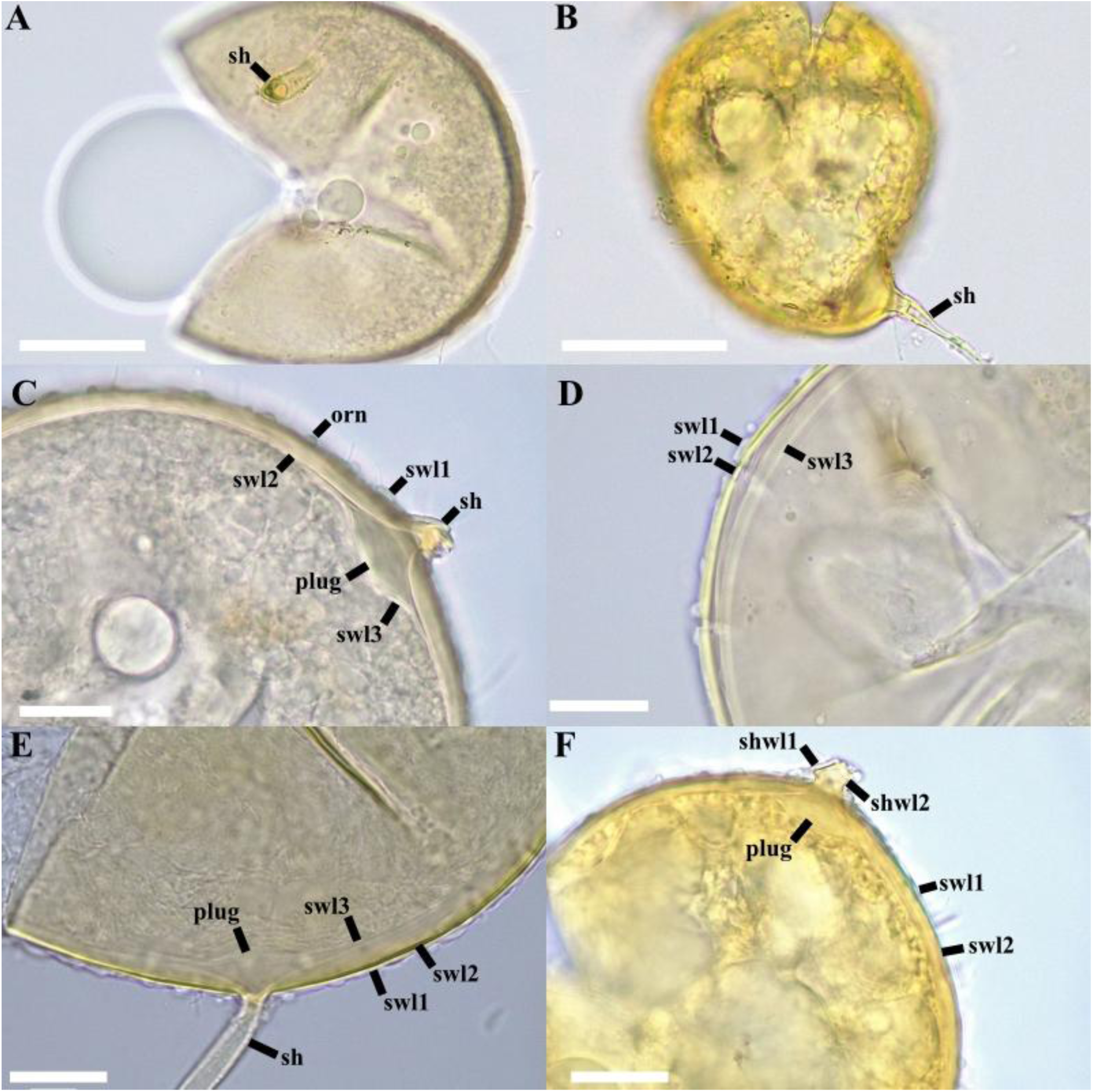
*Diversispora papillosa.* (A, B) Crushed spores in PVLG (A) and PVLG + Melzer (B). (C) ornamentation (orn) and spore wall (swl) 1 – 3, (C-E) Spore wall layers (swl) 1 – 3. (E, F) Subtending hypha (sh) with two wall layers (shwl) 1 – 2. (C, E, F) Plug-like structure. A, C, D, E in PVLG; B, F in PVLG + Melzer’s reagent. Scale bars: (A, B) = 50 μm, (C, D, E, F) = 20 μm.

**MycoBank No:** xxx (will be added after reviews).

**Etymology:** Latin, *papillosa* (= papillae), referring to the surface of the spores, which is adorned with small bumps.

**Specimens examined**: Isolated from rhizospheric soil of a greenhouse pot of a single–species culture propagated on *Sorghum bicolor* at the laboratory Aura Pacifica in New Caledonia, Noumea, June 2024, T. Crossay. This single–species culture was originally inoculated with 100 spores isolated from the rhizospheric soil of *Ficus carica* sampled at a naturally vegetated site located in Fakarava Island (16°08’23.8“S 145°35’26.9”W) in French Polynesia, October 2022, T. Crossay. Holotype deposited at the Mycological Herbarium of the National Museum (France, Paris) in MNHN–PC–(will be added after reviews), isotype (will be added after reviews).

**Diagnosis:** It differs from other species of the genus *Diversispora* in (i) the presence of ornamentation on the spore surface resembling small bulges or bubbles, (ii) the formation of a plug-like structure at the attachment point of the subtending hypha, and (iii) the nucleotide composition of 45S sequences.

**Description**: Glomerocarps unknown. Glomerospores produced singly in soil, blastically at the tip of a sporogenous hypha. Spores globose with one subtending hypha, beige to light grey (6B2) to brownish orange (6C8), translucent, (95–)110(–130) μm diam. (Figure 8A, B, C). Spore wall composed of three layers (Figure 8C-F). Layer 1 (SWL1), forming the spore surface, semi-persistent, ornamented, forming irregular small bulges (1.2–2.2 μm height) on the spore surface (Figure 8C-E), hyaline, (1.8–)2.1(–2.2) μm thick (Figure 8C-E). Layer 2 (SWL2) uniform, smooth, permanent, laminated, light yellow (1A6) (Figure 8D, E), (6.0–)7.1(–8.0) μm thick. Layer 3 (SWL3) uniform (without visible sublayers), flexible, hyaline (Figure 8D, E), (1.2–)1.3(–1.6) μm thick, generally tightly adherent to layer 2. None of the spore wall layers stains in Melzer’s reagent (Figure 8B, F). Subtending hypha hyaline, straight, cylindrical, sometimes constricted at spore base; (3.0–)3.5(–5.0) μm wide at spore base (Figure 8C, E, F). Wall of subtending hypha yellowish white; (2.0–)2.5(–3.3) μm thick at spore base; continuous with layers 1 and 2 of spore wall (Figure 8F). Pore (6.0–)6.4(–7.0) μm in diameter, open or occluded by a plug formed by layers 2 and 3 at the spore base (Figure 8C, E, F). In mounted spores, the formation of a plug-like structure can be observed in the area where the subtending hypha inserts separating layers 2 and 3 (Figure 8C, E, F). Spore content of hyaline oily substance. Germination unknown. Forming hyphal coils and arbuscules, staining with Trypan Blue in cortical roots of *Sorghum bicolor*.

**Distribution and habitat:** In the field, *D. papillosa* has only been reported from the type locality in Fakarava Island, French Polynesia. Only a single match (EUK1187834, 97.2% identity) was found in EUKARYOME v.1.9.4, suggesting a possible occurrence in a tropical broadleaf forest in Puerto Rico.

## Discussion

The incongruencies between the genera *Redeckera* and *Corymbiglomus*, whose species sequences were interspersed within a supported broader clade, were first shown by Tedersoo et al. (2024a), although no formal taxonomic action was taken.

Błaszkowski et al. (2025) “solved” part of the problem, establishing *Paracorymbiglomus* to accommodate *Corymbiglomus globiferum* and *C. pacificum*. In the present study, new species and genera are introduced in the Diversisporales and the polyphyly of *Redeckera* is addressed. Phylogeny was inferred using specimen-based rDNA sequences of Diversisporales and eDNA sequences, providing a broader view of the phylogenetic relationships within the order. Both analyses confirmed the split of *Redeckera sensu* C. Walker and A. Schüßler into three genera, nominally *Redeckera*, *Albocarpum,* and *Pulvinocarpum*. Furthermore, three additional lineages, potentially representing a new family within Diversisporales and two new genera within Corymbiglomeraceae, were also detected in the eDNA phylogeny, but the absence of representative morphospecies precluded their formal recognition. Nonetheless, the use of eDNA sequences from EUKARYOME and their associated metadata proved, once again, to be a powerful approach for identifying candidate novel taxa (Tedersoo et al. 2024b) and for guiding targeted efforts to recover and characterize these organisms.

Notably, among the newly described species, *Albocarpum arenaceum* presented spores in loose aggregates, unlike the glomerocarps characteristic of the congeneric species *A. fulvum* and *A*. *leptohyphum*, as well as all the *Redeckera* species *sensu* C. Walker and A. Schüßler. *Albocarpum arenaceum* culture was, in fact, established using single spores obtained from a trap culture. This unexpected difference in spore formation between species within the same genus reinvigorates the sporulation dimorphism hypothesis proposed by Tedersoo et al. (2024a) to explain the nestedness of *Corymbiglomus* within the *Redeckera sensu* C. Walker and A. Schüßler clade.

Single spores and sporocarps might represent two distinct dispersal strategies, operating locally and distally, respectively. Under this hypothesis, the germination of spores from a sporocarp might depend on a pre-germination process, such as passing through the digestive tract of, for example, a rodent (Jobim et al. 2019a,b; Gil-Fernández et al. 2025). This would prevent the spores from germinating before being dispersed far from the source. The sulfurous odor (egg-like), released by fresh sporocarps of *Albocarpum*, *Melanocarpum*, and *Redeckera* (pers. obs.), would align with this scenario, serving as an attractant for potential vectors. Sporulation dimorphism, therefore, remains a plausible working hypothesis at present, requiring further efforts to obtain isolates in pure culture exhibiting both behaviors, and to verify a potentially concurrent spore dimorphism.

Nevertheless, even in the case of a proven sporulation dimorphism, we reject the option to aggregate under a single genus the species belonging to *Redeckera sensu* C. Walker and A. Schüßler, *Corymbiglomus*, *Paracorymbiglomus*, and the newly introduced *Melanocarpum* since the resulting clade was not convincingly supported (67) in the maximum likelihood phylogeny inferred using representative species of Diversisporales. Furthermore, the percentages of dissimilarity calculated from BLASTn comparisons among sequences of *Corymbiglomus*, *Paracorymbiglomus*, and the genera resulting from the split of *Redeckera* are consistent with values observed in other intergeneric comparisons in the Glomeromycota (e.g., *Albahypha* vs *Entrophospora*: 9.5%; *Racocetra* vs *Cetraspora*: 6.1%; *Dominikia* vs *Macrodominikia*: 9.3%).

Among the *Redeckera sensu* Oehl et al. (2011a) species, *R*. *avelingiae*, *R*. *canadensis*, and *R*. *fragilis* were not included in the analysis. Due to the lack of molecular data, these three species remain as *Redeckera*, to avoid proliferation of names, pending additional morphological and phylogenetic analysis.

*Redeckera varelae* and *R. megalocarpa* share a moderately high percentage of identity (96%). However, they differ in size and shape of the glomerocarps. In *R. megalocarpa*, glomerocarps are three times larger (38 × 20 mm) and irregularly shaped, whereas *R. varelae* forms compact, pulvinate glomerocarps, with 14 × 5 mm diam. The most important difference between *R. megalocarpa* and *R. varelae* is the composition of the spore wall. The former species presents only two spore wall layers, whereas *R. varelae* presents three spore wall layers. In *R. varelae*, the wall is up to 1.1 μm thicker than in *R*. *megalocarpa*. In addition, the laminated layer of the *R. varelae* stains pink in Melzer’s reagent (14A3), whereas in *R. megalocarpa* any layer presents a Melzer’s reaction. Originally, *R*. *megalocarpa* (=*G*. *megalocarpum*) was described as having a single-layered wall (Redecker et al. 2007). However, the picture 109 in Oehl et al. (2011a) clearly shows two wall layers in this species.

In *Albocarpum*, *A*. *leptohyphum* was preferred as the designated type species rather than *A. fulvum*, despite the latter having been described more than a century and a half earlier (Berkeley and Broome 1873), due to the absence of fresh material from the type location (Sri Lanka) and a detailed morphological description. *Albocarpum arenaceum* differs significantly from the other two species in the genus for having a different mode of spore formation, i.e., it produces loose aggregates with 5–15 spores in single species culture, while *A*. *leptohyphum* and *A*. *fulvum* form large sporocarps in the field covered by a cottony-looking peridium and containing thousands of spores within the gleba. Apart from this aspect, *A*. *arenaceum* and *A*. *leptohyphum* share globose to subglobose spores of similar diameter (70–120 μm), and a three-layered wall. However, they differ mainly in the morphology of the spore wall and the subtending hypha. In *A*. *arenaceum*, the spore wall is slightly thicker (up to 1.1 μm) than in *A*. *leptohyphum*. Additionally, the outermost layer of the former species is mucilaginous, whereas in the latter species is persistent and semi-flexible, easy to detect even in mature spores. Another diagnostic difference concerns the septum occluding the subtending hypha: in *A*. *leptohyphum* the septum is straight and positioned at the base of the spore, whereas in *A*. *arenaceum* it is curved and formed up to 15 µm below the spore base. Both species are easily distinguishable from *A*. *fulvum*, which produces ellipsoid to ovoid spores with a three-layered wall (Thaxter 1922; Oehl et al. 2011a; Błaszkowski 2012).

*Pulvinocarpum pulvinatum*, like the other species previously assigned to *Redeckera sensu* C. Walker and A. Schüßler, forms large sporocarps, a feature representing a synapomorphy also observed in several genera of the Glomerales, including *Epigeocarpum*, *Dominikia*, and *Sclerocarpum* (Jobim et al. 2019a; Błaszkowski et al. 2021). Consequently, sporocarp formation alone cannot be considered a reliable diagnostic character without detailed evaluation of the phenotypic and histochemical properties of the spore wall and the subtending hypha (Oehl et al. 2011a; Goto et al. 2024). The main differences between *P. pulvinatum* and *Redeckera* species are the formation of globose to subglobose spores with a three-layered wall, non-reactive to Melzer’s reagent, and the nucleotide composition of 45S barcode sequences (Thaxter 1922; Gerdemann and Trappe 1974; Redecker et al. 2007; Oehl et al. 2011a; Błaszkowski 2012). Morphologically, the species with which *P*. *pulvinatum* could be confused is *A*. *leptohyphum* due to the size and shape of the sporocarp and spores. However, the spores of *P*. *pulvinatum* are globose to subglobose (Gerdemann and Trappe 1974) and slightly smaller than those of *A*. *leptohyphum*. In addition, the laminated layer of *A*. *leptohyphum* is up to three times thicker than that of *A. pulvinatum* (Gerdemann and Trappe 1974).

The morphological characteristics of *Melanocarpum mexicanum* resemble, at first, those of *Diversispora epigaea* and *D*. *sporocarpia* (Daniels and Trappe 1979; Jobim et al. 2019a), the only two species within *Diversispora* that form large, compact, unorganized epigeous sporocarps, covered by a peridium and containing hundreds of spores. However, the differences in the 45S sequences and the phylogenetic placement undoubtedly separate *M. mexicanum* from members of *Diversispora*. Although all three species produce sporocarps of similar shape and color, those of *D. sporocarpia* are the smallest (6 × 4 mm) compared to *D. epigaea* (2–8 × 3–15 mm), and *M*. *mexicanum* (8–15 × 4–7 mm) (Daniels and Trappe 1979; Jobim et al. 2019a). Additionally, the spores of *M*. *mexicanum* (250 µm diam.) are on average twice as large as those of *D*. *epigaea* and *D*. *sporocarpia* (129 and 136 µm diam., respectively). The spore wall of *M. mexicanum* and *D. sporocarpia* has three and four layers, respectively. In both species, the first layer exhibits a similar phenotype; however, when intact, it is slightly thinner in *M. mexicanum*. In *D. sporocarpia*, the second layer is uniform and without visible sublayers, whereas in *M. mexicanum* is thick, finely laminated (20 μm) with a fibrous appearance and irregular surface, as in the interior of the second layer of *Funneliformis kerguelense* (see Figure 2G in Dalpé et al. 2002). Another difference between the two species lies in the third layer, which in *D. sporocarpia* is flexible, permanent, finely laminated, and relatively thick (5–9.8 µm). This layer is followed by a fourth, uniform, thin (0.4–0.7 µm), permanent, and flexible layer. In *M. mexicanum*, by contrast, the third layer lacks distinct sublayers, is permanent, flexible, and much thinner (0.5–1.5 µm). The differences between the spore wall of *M. mexicanum* and *D. epigaea* are very clear; in the former species, the wall is composed of three layers and is 25 µm thick, whereas the wall of *D. epigaea* has two layers and is up to 10 µm thick (Daniels and Trappe 1979). Finally, the second layer in *M. mexicanum* stains dark orange (6B8) in Melzer’s reagent, whereas none of the layers in *D. sporocarpia* or *D. epigaea* presents a reaction.

Analysis of partial 45S sequences revealed the formation of a fully supported clade composed of *Diversispora trimurales*, *D*. *gibbosa*, *D*. *peridiata*, and the new species described here as *D*. *papillosa*, sister to the clade of other *Diversispora* species. In spite of the molecular divergences, we resigned from the temptation of establishing a new genus as we did not detect clear morphological features separating the two clades. Of the four species, *D*. *papillosa* and *D*. *trimurales* form solitary spores only, while *D. gibbosa* forms both solitary spores and loose aggregates enclosed by a hyphal mantle. In contrast, *D*. *peridiata* forms clusters of 3–20 spores covered by a peridium, and has the smallest globose spores, while the other three species show similar spore diameter (110–128 μm) (Koske and Halvorson 1989; Błaszkowski 2012; Błaszkowski et al. 2015). The wall structure of *D. papillosa* differs significantly from its three sister species (*D. gibbosa*, *D*. *trimurales*, and *D*. *peridiata*). The main difference resides in the SWL1, which is evanescent and smooth in *D. gibbosa* (Błaszkowski 1997; 2012), whereas in *D*. *papillosa* it is semi-persistent and ornamented with small protuberances. The spore wall of *D*. *trimurales* and *D. papillosa* is adorned with protuberances; however, their height is lower in *D*. *papillosa* (1.2–2.1 μm vs. 3.1–5.9 μm high). Furthermore, in PVLG, the SWL2 of *D*. *trimurales* separates from SWL3 (Koske and Halvorson 1989; Błaszkowski 2012), whereas in *D*. *papillosa*, layers 2 and 3 remain strongly adhered even when pressure is applied.

Environmental sequences from EUKARYOME v.1.9.4 (Tedersoo et al. 2024a) not only supported the reorganization of the *Redeckera*–*Corymbiglomus*–*Paracorymbiglomus* group but also enabled the mapping of the global distributions for the newly established genera and species. Interestingly, most occurrences were recorded in the Pantropical zone, which has been shown to be a hotspot of AMF diversity (Van Nuland et al. 2025). *Albocarpum* showed a broad distribution across most biomes of this bioregion, whereas *Pulvinocarpum*, *Redeckera*, and *Melanocarpum* were mainly restricted to tropical biomes of the Americas. Notably, occurrences of *Melanocarpum* suggested a possible endemism confined to the Antilles region. Among the *Albocarpum* species, *A. arenaceum* showed the broadest distribution, while other species displayed more limited geographic ranges. The occurrences of *A. fulvum*, including the specimen from Martinique whose partial 45S sequences were used by Redecker et al. (2007) to define the lineage, indicated a possible endemism in Caribbean tropical forests, even though the specimen used to describe the species (as *Paurocotylis fulva*) was collected in Sri Lanka (Berkeley and Broome 1873). Similarly, *R. varelae* was detected in tropical forests of two Caribbean countries, other than the sampling site of the specimen. A more strict endemism was shown for *M. mexicanum*, as the sequences potentially related to the species were obtained from different sites, all in Mexico. Finally, *A. leptohyphum* and *D. papillosa* appeared to be very rare species, the former lacking any related environmental sequence, and the latter represented by only a single environmental sequence found in EUKARYOME. It should be emphasized that the reliability of these patterns depends strongly on the number and geographic distribution of sequenced samples. As the EUKARYOME database continues to expand with long-read Glomeromycota sequences, clearer and more robust distributional trends are expected to emerge.

## Conclusions

Five new species and a new genus in the Diversisporales are described, together with the split of *Redeckera* into three genera. In addition, an epitype was designated for *A*. *fulvum* (former *R. fulva*), and its description was emended accordingly, supported by isolate-and eDNA-based phylogenies. Remarkably, three of the five novel species were characterized from glomerocarpic specimens collected in the field, a leap to the earliest period of Glomeromycota discovery when the description of the first species relied on sporocarp-forming fungi (Gedermann and Trappe 1974). These results assume even greater significance considering that, over the past decade, the description of glomerocarpic species *sensu stricto* (Jobim et al. 2019a) has been limited to only six species (Jobim et al. 2019a, Blaszkowski et al. 2019, 2021, Yamato et al. 2024). This pattern is unlikely to reflect true rarity, but rather the difficulties associated with detection, characterization, cultivation, and, last but not least, the possibility that large sporocarps represent a peculiar mode of sporulation, alternative to single spore formation, and strongly influenced by seasonality, edaphic, and vegetation conditions.

## Data availability statement

The datasets presented in this study are available in online repositories. The names of the repository/repositories and accession number(s) are listed below: NCBI PX612380-PX612384; PX661504-PX661521; PX700901-PX700907.

## Acknowledgments

We thank the project of the Tecnológico Nacional de México (23336.25-P) assigned to MHPM and LALP. We thank the Conselho Nacional de Desenvolvimento Científico e Tecnológico (CNPq) for the research grant awarded to B.T.G. (proc.409181/2024-2; 306632/2022-5) and J.R.L.L. (proc.151661/2024-3).

We respectfully acknowledge the late Christine Wong (DAG) for her contributions to this project.

We equally thank Elena Gorchakova (UICN) for her contributions.

